# sFlt-1 commutes unfolded protein response into endoplasmic reticulum stress in trophoblast cells in preeclamptic pregnancies

**DOI:** 10.1101/297374

**Authors:** Sankat Mochan, Manoj Kumar Dhingra, Sunil Kumar Gupta, Shobhit saxena, Pallavi Arora, Neerja Rani, Arundhati Sharma, Kalpana Luthra, Sadanand Dwivedi, Neerja Bhatla, Rani Kumar, Renu Dhingra

**Author notes:** Correspondence: Renu Dhingra.

## Abstract

Preeclampsia (PE) and its subtypes (early and late onset) are serious concerns all across the globe affecting about 8% of total pregnancies and accounts for approximately 60,000 deaths annually with a predominance in developing under-developed and countries. The two-stage model in the progression of this disease, deficient spiral artery remodelling and an imbalance between angiogenic (VEGF) and anti-antigenic factor(s) (sFlt-1) are well established facts pertaining to this disease. The presence of increased sFlt-1, high oxidative stress and Endoplasmic reticulum stress (ER stress) have been proposed in preeclamptic pregnancies. Recently, the role of endoplasmic reticulum stress in the onset of the variant forms of PE highlighted a new window to explore further. In our previous studies, we demonstrated that sFlt-1 can induce apoptosis and oxidative stress in trophoblast cells. However the role of sFlt-1, in inducing ER stress is not known so far. In the present study, we for the first time demonstrated significant ER stress in the placental cells (BeWo Cells) (*in vitro*) when exposed to sera from preeclamptic pregnancies having increased concentration of sFlt-1. The expression of ER stress markers (GRP78, eIF2α, XBP1, ATF6 and CHOP) at both transcript and protein levels were compared (between preeclamptic and normotensive non-proteinuric women) at three different time points (8h, 14h and 24hrs), analyzed and found to be significant (p<0.05).

**Conclusion:** Our results suggested that sFlt-1, released from placental cells in preeclampsia may be one of the various factors having potential to induce endoplasmic reticulum stress in BeWo cells.

## Introduction

sFlt-1, a splice variant of VEGF receptor Flt-1, lacks the trans-membrane and cytoplasmic domains, is made in large amounts by placental trophoblasts and is released into maternal circulation **^1,2^**. It acts as a potent anti-angiogenic molecule by binding circulating VEGF and PlGF. Several studies worldwide reported that sFlt-1 gene product is up-regulated in the preeclamptic placentae**^3,4,5^**. Alterations in the angiogenic molecules (increased sFlt-1, decreased free VEGF and PlGF) antedate the onset of clinical symptoms of preeclampsia (PE) which is generally defined as new hypertension (diastolic blood pressure of >90 mm Hg or systolic blood pressure of >140mm Hg) and substantial proteinuria (≥300 mg in 24 h) at or after 20 weeks of gestation**^6,7^**. It is a common disorder (5-8% of all pregnancies) and often requires premature delivery of the baby**^8,9^**. Occasionally, PE may lead to seizures and multi-organ dysfunction in the mother with severe morbidity and even mortality**^10^**. It occurs only in presence of placenta and remits dramatically postpartum, after the placenta has been delivered**^11^**. The placentae of preeclamptic women appear wayward, with evidence of underperfusion and ischemia**^12^**. The initial event in PE has been posited to be reduced uteroplacental perfusion which develops as a result of abnormal cytotrophoblast invasion of spiral arterioles, triggers the cascade of events leading to the maternal disorder**^13^**. Such malperfusion leads to poor placentation which causes both oxidative and endoplasmic reticulum stress in the placenta**^14^**. The Endoplasmic reticulum (ER) performs multiple functions, including the synthesis, post-translational modification, and trafficking of membrane and secreted proteins**^15^**. Fuss of ER homeostasis may result in misfolding or abnormal glycosylation of these proteins, which may reduce their biological activity**^16^**. The accumulation of unfolded or misfolded proteins within the ER cisternae incites ER stress and activation of the unfolded protein response (UPR) **^17^**. The UPR ventures to restore ER function by attenuating protein translation, increasing folding capacity, and facilitating mortification of misfolded proteins**^18^**. It consist of three leading signaling pathways with overlapping functions**^19^**. The three trans-membrane sensors of UPR consist of PKR-like endoplasmic reticulum kinase (PERK), activating transcription factor 6 (ATF6), and inositol-requiring 1 (Ire1), project into the ER lumen**^20^**. Although several studies have indicated the importance of ER stress in the development of PE but the impact of sFlt-1 on the overall expression profiles of ER stress markers (GRP78, eIF2α, XBP1, ATF6 and CHOP) in BeWo cells has not been reported so far. The present study was designed to explore the effect of sFlt-1 on ER stress in the trophoblast cells (*in vitro* study). We demonstrated ER stress in BeWo cells (human choriocarcinoma cell line of placental origin) because it mimics *in vivo* syncytialisation of placental villous trophoblast.

## Materials and Methods

### Study Subjects

60 pregnant women were enrolled from the antenatal clinic and the inpatient ward of the Department of Obstetrics and Gynaecology, All India Institute of Medical Science, New Delhi, India. The preeclamptic women (n=30) after clinical diagnosis were enrolled as cases and normotensive, non-proteinuric pregnant women (n=30) (maternal and gestational age matched) without any other medical complications were enrolled as controls. Protocol of the study was approved by the institute ethics committee and written informed consent was obtained from all the enrolled women. Preeclampsia was defined according to ACOG guidelines: Blood Pressure-140 mm Hg systolic or >90mm Hg diastolic on 2 occasions at least 4 hours apart after 20 weeks of gestational age in women with a previously normal BP and 160 mm Hg systolic or >110 mm Hg diastolic, confirmed within a short interval (minutes) to facilitate timely antihypertensive therapy; Proteinuria >300 mg per 24-hour urine collection or protein/ creatinine ratio > 0.3mg/dl or dipstick reading of >1+ or in the absence of proteinuria, new-onset hypertension with new onset of one or more of the following thrombocytopenia: platelet count <100,000/μl, renal insufficiency: serum creatinine > 1.1mg/dl or doubling of serum creatinine in the absence of other renal disease, impaired liver function: elevated blood levels of liver transaminases to twice normal concentrations, pulmonary edema and cerebral edema. Pregnant women with chronic hypertension, chorioamnionitis, diabetes, renal disease, cardiac disease were excluded from the study. 5 ml of venous blood was collected, centrifuged at 1200 RPM for 4 minutes, serum was separated and stored in aliquots at -80°C. The serum samples were stored for the ELISA and cell culture experiments.

### Serum analysis of sFlt-1, VEGF and GRP78 using ELISA

The levels of sFlt-1,VEGF and GRP78 were estimated in the serum of preeclamptic and normotensive, non-proteinuric pregnant women groups by sandwich ELISA (sFlt-1 and VEGF ELISA kit: R&D Systems Inc., Minneapolis, MN, U.S.A., GRP 78: Enzo Life Sciences, Inc.).

***In vitro* experiments** were carried out to analyze the effect of sFlt1-1 on induction of ER stress in trophoblastic cells (BeWo cells). The human choriocarcinoma cell line (BeWo) was procured from American Type Culture Collection (ATCC) and maintained in F-12 HAM nutrient medium supplemented with 10% fetal bovine serum, 100 U/ml pencillin, 100μg/ml Streptomycin. Cells were passaged with 0.025% trypsin and 0.01% EDTA.

The study was divided into seven experimental groups depending on various treatments given to BeWo cells: group 1 [normotensive, non-proteinuric serum (low concentration of sFlt-1)], group 2 [normotensive, non-proteinuric serum along with recombinant sFlt-1 (NT+re-sFlt-1)], group 3 [preeclamptic serum (high concentration of sFlt-1)], group 4 [preeclamptic serum along with recombinant VEGF (PE+re-VEGF)], group 5 [recombinant sFlt-1 (re-sFlt-1)], group 6 (tunicamycin treatment), group 7 (untreated)]. After the various treatments, activation of ER stress markers (GRP78, eIF2α, XBP1, ATF6 and CHOP) were assessed at various time points (8h, 14h, 24h) at protein level (Immunofluorescence, Western blot) and gene level (qRT-PCR).

### Immunofluorescence Microscopy

BeWo Cells were trypsinized, seeded, allowed to grow on coverslips in multiple well chamber and incubated at 37°C in 5% CO2. After 8h, 14 h and 24 h, cells were taken out from incubator and washed with PBS, fixed in 4% PFA for 15 min at room temperature. After fixation, cells were washed with PBS and permeabilized with PBS + 0.1% Triton X-100 followed by PBS washing. Nonspecific blocking was done using 5% normal goat serum in PBS and Triton X. Cells were incubated for 12 hours at 4°C in primary antibodies (anti GRP78 antibody (1:1000), anti eIF2α antibody (1:200), anti XBP1 antibody (1:200), anti ATF6 antibody (1:1000) and anti DDIT3/CHOP antibody (1:500). Cells were washed with PBSTx and thereafter incubated in secondary antibody in 1:500 dilution for 1 hour at room temperature in dark room. Cells were washed in PBS and mounted with flouroshield mounting media with DAPI on the slide and observed under the fluorescence microscope (Nikon Eclipse Ti-S elements using NiS-AR software).

### Western blot analysis

Cells were lysed in SDS-PAGE sample buffer [2% SDS, 60 mM Tris-HCl (pH 6.8), 10% glycerol, 0.001% bromophenol blue, 0.33% mercaptoethanol] and boiled for 5 min. The lysates were analyzed by immunoblotting using 1:1000 of anti GRP78, 1:500 of anti eIF2α, anti XBP1, and 1:1000 of anti ATF6, anti CHOP (Abcam) for 12 h at 4°C. The blots were then incubated in secondary antibody (HRP conjugated) for 2 hours. The membranes were treated with DAB. β-actin was used as protein loading control. The immunoreactive proteins were visualized using the gel documentation system (Bio-Rad, Hercules, CA, USA, Quantity 1 software).

### qRT-PCR (quantitative Real Time-Polymerase Chain Reaction)

RNA extraction from the treated cells was done using Ambion (Invitrogen) kit followed by c-DNA conversion by Thermo revert aid H-minus reverse transcriptase kit. cDNA was amplified by quantitative RT-PCR for determining mRNA expression of GRP78, eIF2α, XBP1, ATF6 and CHOP against gene of interest with an internal control (β actin and GAPDH).

### Statistical Analysis

Data was analyzed by Microsoft Office Excel Version 2013 and Graph Pad Prism. Relative quantification cycles of gene of interest (ΔCq) was calculated by ΔCq = Cq (target) - Cq (reference). Relative mRNA expression with respect to internal control gene was calculated by 2^-ΔCq^. The data are presented as mean±SD for VEGF, GRP78 and median for sFlt-1. Average level of the variable between the two groups was compared by paired t-test/Wilcoxon signed rank test. For comparing more than two groups, ANOVA with Bonferroni’s multiple comparison test/Kruskal Wallis with Dunn’s multiple comparison test were used. *p* value<0.05 was considered statistically significant.

## Results

Maternal serum of 60 pregnant women were analyzed. The mean systolic and diastolic blood pressures in preeclamptic group were 158.9 ± 11.88 mm Hg (mean ± SD) and 101.43 ± 8.39 mm Hg (mean ± SD) respectively where as in control group, the systolic and diastolic pressures were 117.8 ± 7.34 mm Hg (mean ± SD) and 74.2 ± 6.39 mm Hg (mean ± SD) respectively. The difference in the systolic and diastolic pressures between the groups were statistically significant (*p*<0.0001). The body mass index in preeclamptic patients was 27.83± 5.61 (mean ± SD) and in control group was 23.67 ± 3.59 (mean ± SD). The difference was statistically significant (*p*<0.0001). Urine protein in preeclamptic and control group women were analyzed by urine dipstick method. 10% of preeclamptic women (3/30) showed 1+urine protein, 43.33% showed 2+ urine protein (13/30) and 46.66% showed 3+ urine protein (14/30) (Table 2)

### Elevated levels of sFlt-1 and GRP-78 and reduced VEGF levels were observed in preeclamptic patients sera

The level of sFlt-1 in the maternal serum of preeclamptic patients was 11295.25 (2936.2-37818) pg/ml and in control group was 2936.2 (1180.43-6706.6) pg/ml. The serum sFlt-1 levels were significantly higher in women with preeclampsia than in control group and the difference was statistically significant. (*p*=0.0001) (Table 3, Figure 2). The level of GRP78 in the maternal serum of preeclamptic patients was 1103.26 ± 104.27 ng/ml and in control group was 1018.61 ± 125.51 ng/ml. The serum GRP 78 levels were higher in sera of women with preeclampsia than in control group and the difference was statistically significant. (*p*=0.012) (Table 3, Figure 3). Patients with preeclampsia showed reduced serum levels of VEGF 170.53 ± 36.55 pg/ml (mean± SD) than normotensive non-proteinuric pregnant women 254.61 ± 47.39 pg/ml (mean± SD) (p< 0.0001) (Table 3, Figure 4).

**Figure 1.**
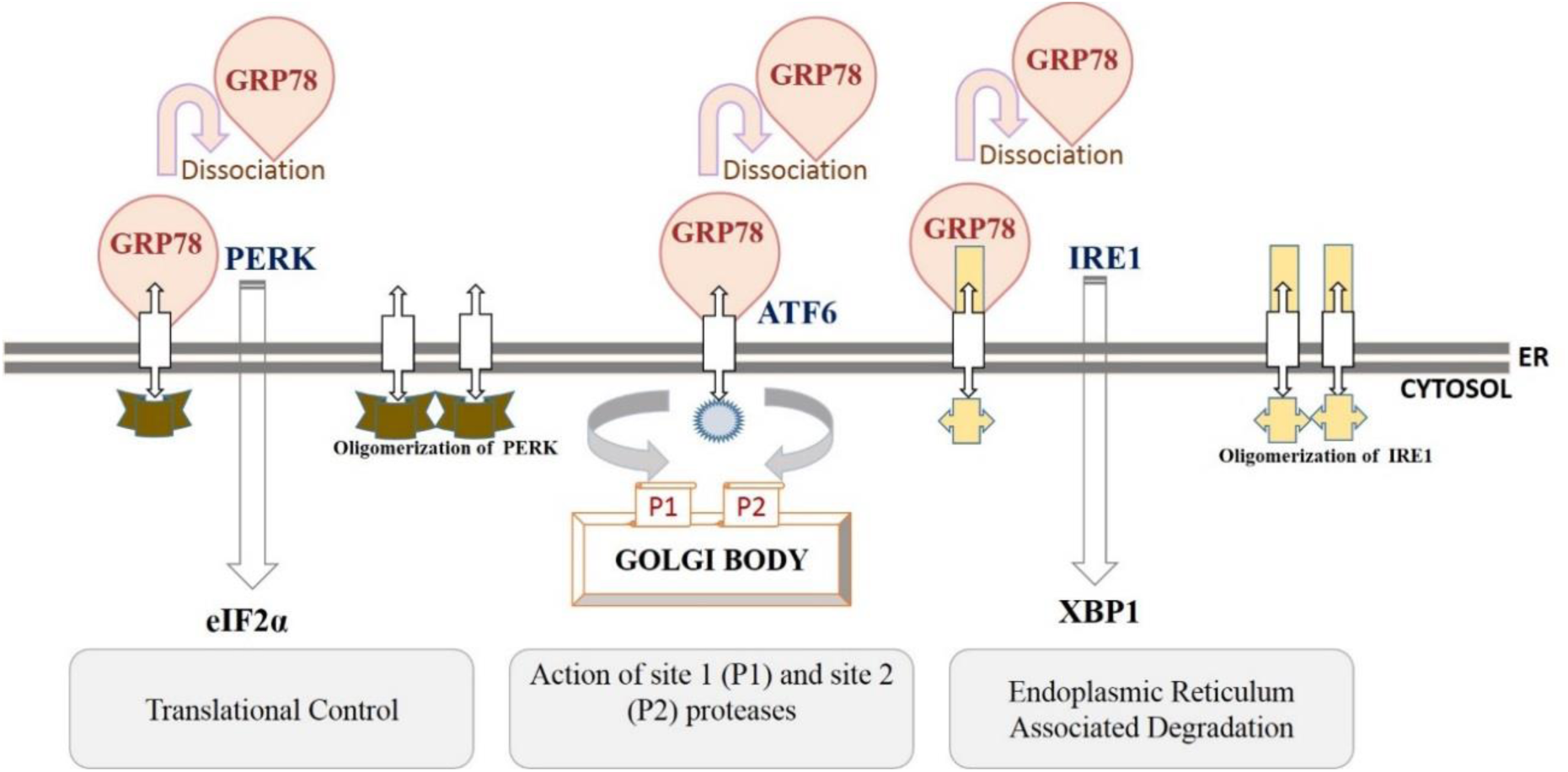
Endoplasmic reticulum stress pathways

**Figure 2:**
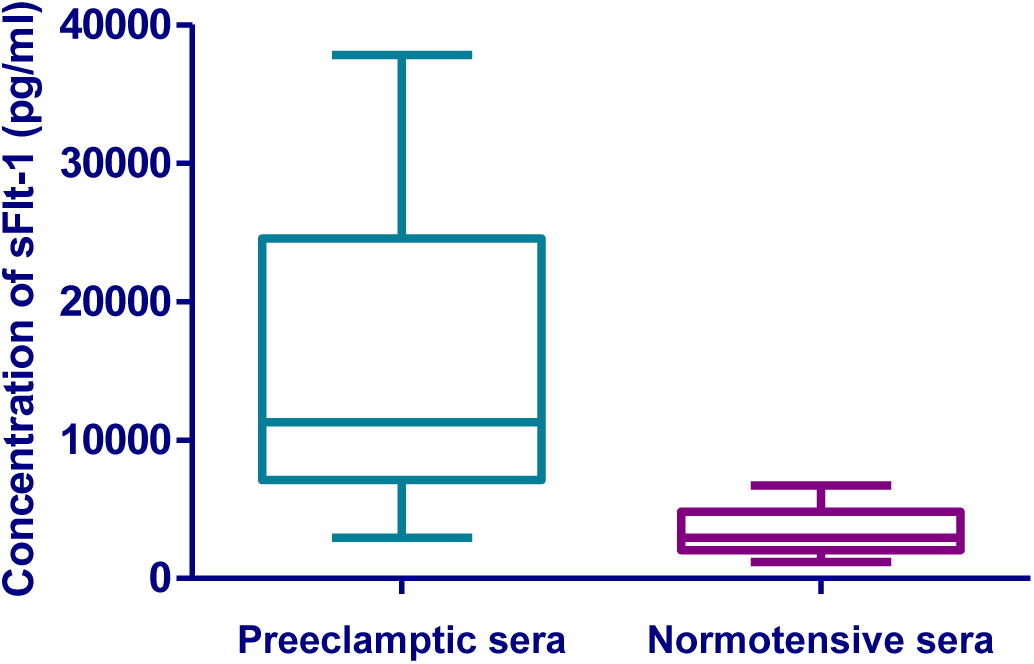
Box plots showing serum levels of soluble fms-like tyrosine kinase-1 (sFlt-1). Boxes denote the interquartile range with the upper and lower horizontal edges representing the 75^th^ and 25^th^ percentiles. The central horizontal lines represent the medians. *Statistical significance, *p*<0.05

**Figure 3:**
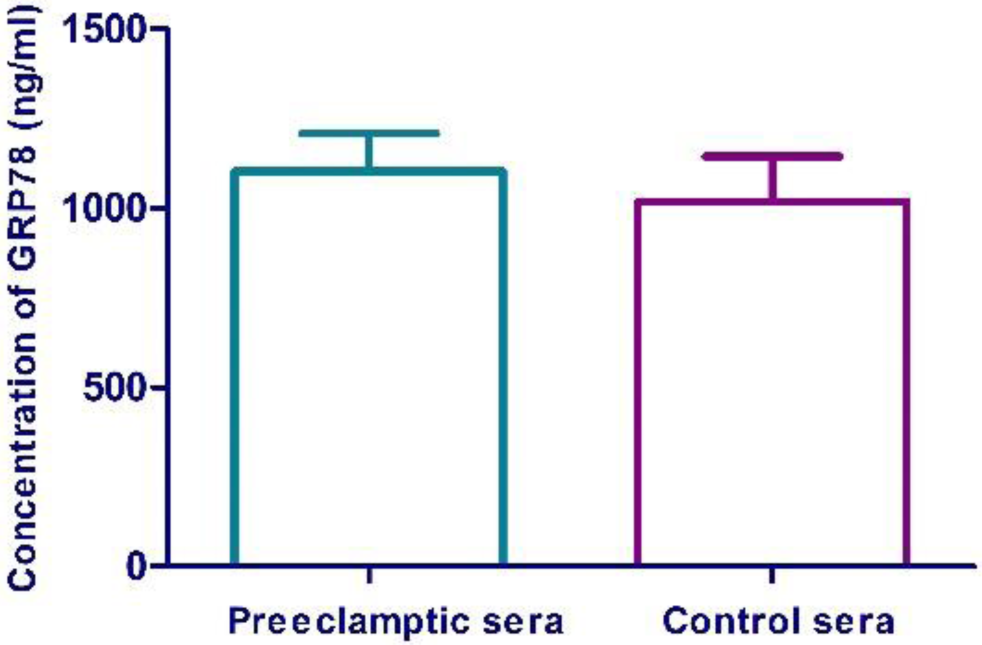
Bar diagram represents serum levels of Glucose Regulated Protein78 (GRP78). Values are represented as Mean +SD. Error bars represent standard deviation. Statistical significance, *p*<0.05

**Figure 4:**
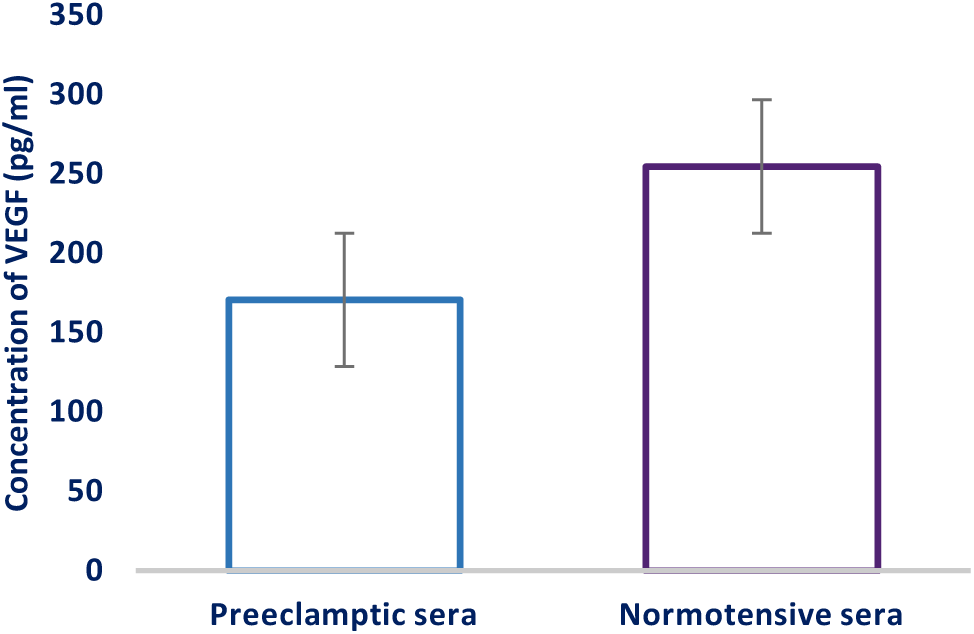
Bar diagram represents serum levels of Vascular VEGF. Values are represented as Mean +SD. Error bars represent standard deviation. Statistical significance, *p*<0.05

### *In Vitro* Experiments

#### Weak expression of ER stress markers was observed in BeWo cells treated with sera from Normotensive (NT) pregnant women [Figure 5a-c]

Immunofluorescence microscopy revealed no significant expression of ER stress markers at different time points [Figure 5a] nevertheless immunoblot manifested higher expression of GRP78 and eIF2α at 14 h as compared to 8 h and 24 h whereas XBP1, ATF6 and CHOP expressions were found more at 24 h as compared to 8h and 14 h [Figure 5b]. mRNA levels of GRP78, eIF2α, XBP1, ATF6 and CHOP were found to be higher at 14 h as compared to 8 h and 24 h [Figure 5c].

**Figure 5a:**
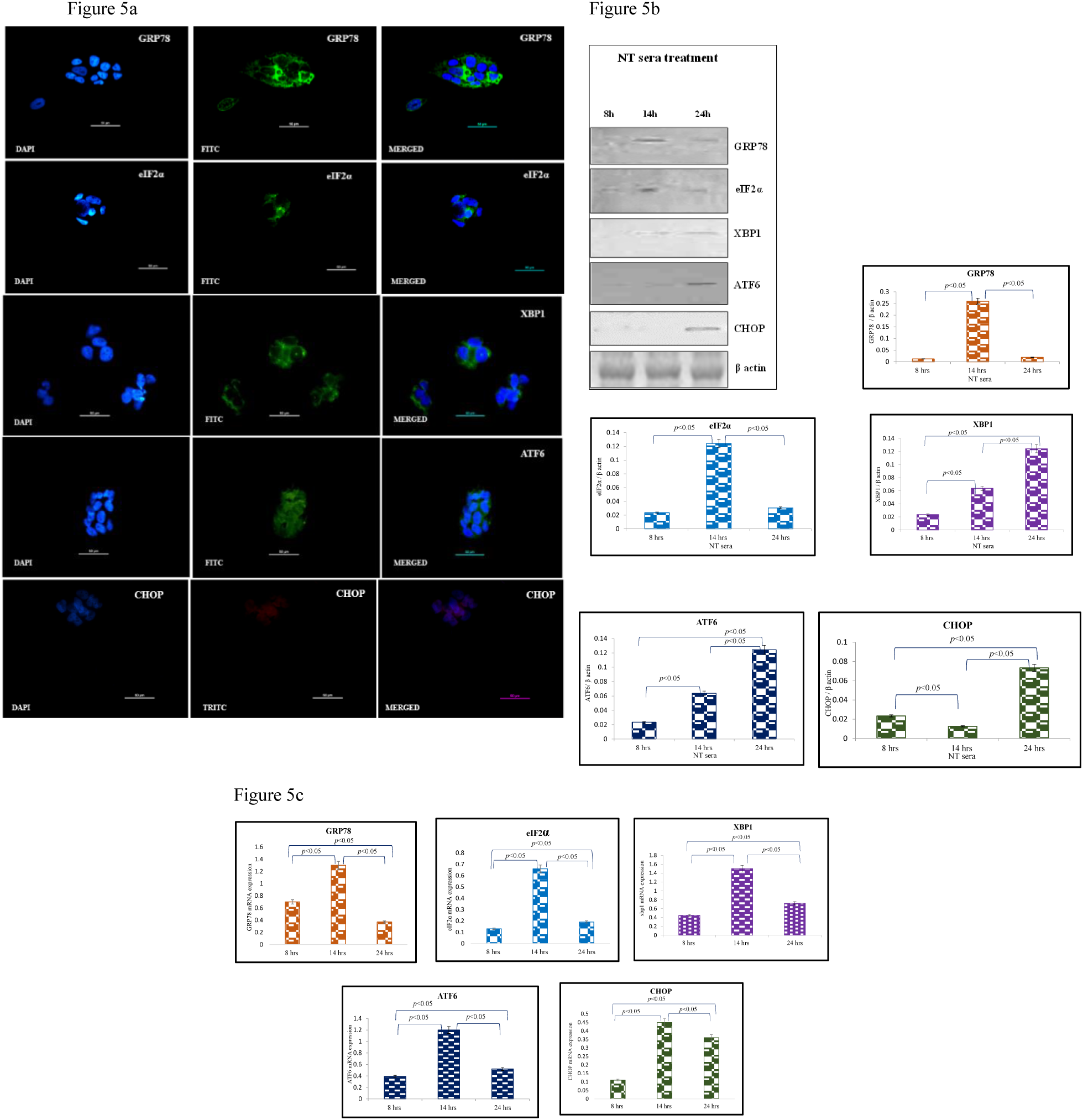
Representative immunofluorescence staining pattern of anti-GRP78 antibody positive BeWo cells following 8 h, anti-eIF2α, anti-XBP1, anti-ATF6 antibody positive BeWo cells following 14 h and anti-DDIT3 (CHOP) antibody positive BeWo cells following 24 h treatment of NT sera; Figure 5b: Representative images of immunoblot showing the expression of ER stress markers, GRP78, eIF2α, XBP1, ATF6 and CHOP in BeWo cells. β-Actin was used as protein loading control. The Bar diagrams represent the normalized values of the markers. Results are representative of 7 independent experiments. Data presented as mean ± SD. Statistical analysis was done using one way ANOVA with Bonferroni’s post hoc; Figure 5c: Bar diagrams represent the relative mRNA expression of GRP78, eIF2α, XBP1, ATF6 and CHOP and was found maximum at 14 h. GAPDH was used as positive control. Data presented as mean ± SD. One way ANOVA with Bonferroni’s post hoc test was applied (*p* values indicated on graph itself)

#### Expression of ER stress markers in BeWo cells was enhanced on treatment with the recombinant sFlt-1(re-sFlt-1) added to sera from normotensive pregnant women [Figure 6a-c]

Immunofluorescence microscopy revealed significant expression of GRP 78 at 8 hours, eIF2α, XBP1, ATF6 at 14 hours and CHOP at 24 hours. Characteristic stress granules in case of eIF2α were observed at 14 hours [Figure 6a]. The expression of GRP78, eIF2α and XBP1 proteins were found to be higher at 14 h as compared to 8 h and 24 h time points. ATF6 expression was found to be higher at 8 and 24 h time points as compared to 14 h. Maximum expression of CHOP was found at 24 h [Figure 6b]. mRNA levels of GRP78, eIF2α and XBP1 were found to be higher at 14 h whereas ATF6 and CHOP mRNA levels were found to be higher at 24 h [Figure 6c].

**Figure 6a:**
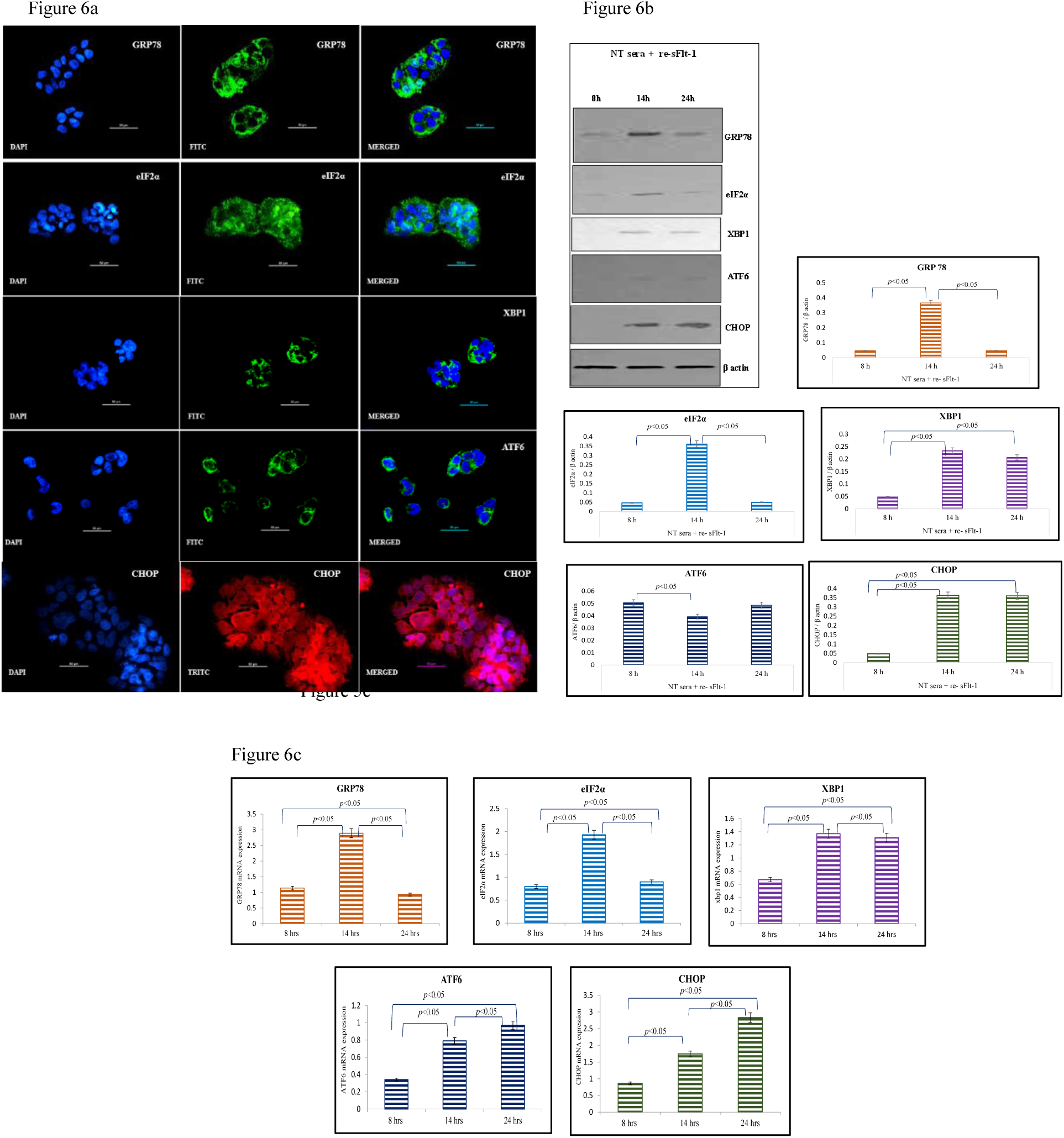
Representative immunofluorescence staining pattern of anti-GRP78 antibody positive BeWo cells following 8 h, anti-eIF2α, anti-XBP1, anti-ATF6 antibody positive BeWo cells following 14 h and anti-DDIT3 (CHOP) antibody positive BeWo cells following 24 h treatment of NT sera + re-sFlt-1; Figure 6b: Representative images of immunoblot showing the expression of ER stress markers, GRP78, eIF2α, XBP1, ATF6 and CHOP in BeWo cells. β-Actin was used as protein loading control. The Bar diagrams represent the normalized values of the markers. Results are representative of 7 independent experiments. Data presented as mean ± SD. Statistical analysis was done using one way ANOVA with Bonferroni’s post hoc; Figure 6c: Bar diagrams represent the relative mRNA expression of GRP78, eIF2α, XBP1, ATF6 and CHOP. GAPDH was used as positive control. Data presented as mean ± SD. One way ANOVA with Bonferroni’s post hoc test was applied (*p* values indicated on graph itself)

#### Significantly increased expression of GRP78, eIF2α, XBP1, ATF6 and CHOP in Preeclamptic sera (PE) treated BeWo cells was observed [Figure 7a-c]

Immunofluorescence microscopy revealed significant expression of GRP78 at 8 hours, eIF2α, XBP1, ATF6 at 14 hours and CHOP at 24 hours [Figure 7a]. Immunoblot data revealed that expression of GRP78 was found to be higher at 8 h as compared to 14 h and 24 h time points, whereas eIF2α, ATF6 and CHOP protein expressions were found to be higher at 24 h as compared to 8 h and 14 h. Expression of XBP1 was found maximum at 14 h [Figure 7b]. mRNA levels of GRP78, eIF2α, XBP1, ATF6 and CHOP were found to be higher at 14 h as compared to 8 h and 24 h [Figure 7c].

**Figure 7a:**
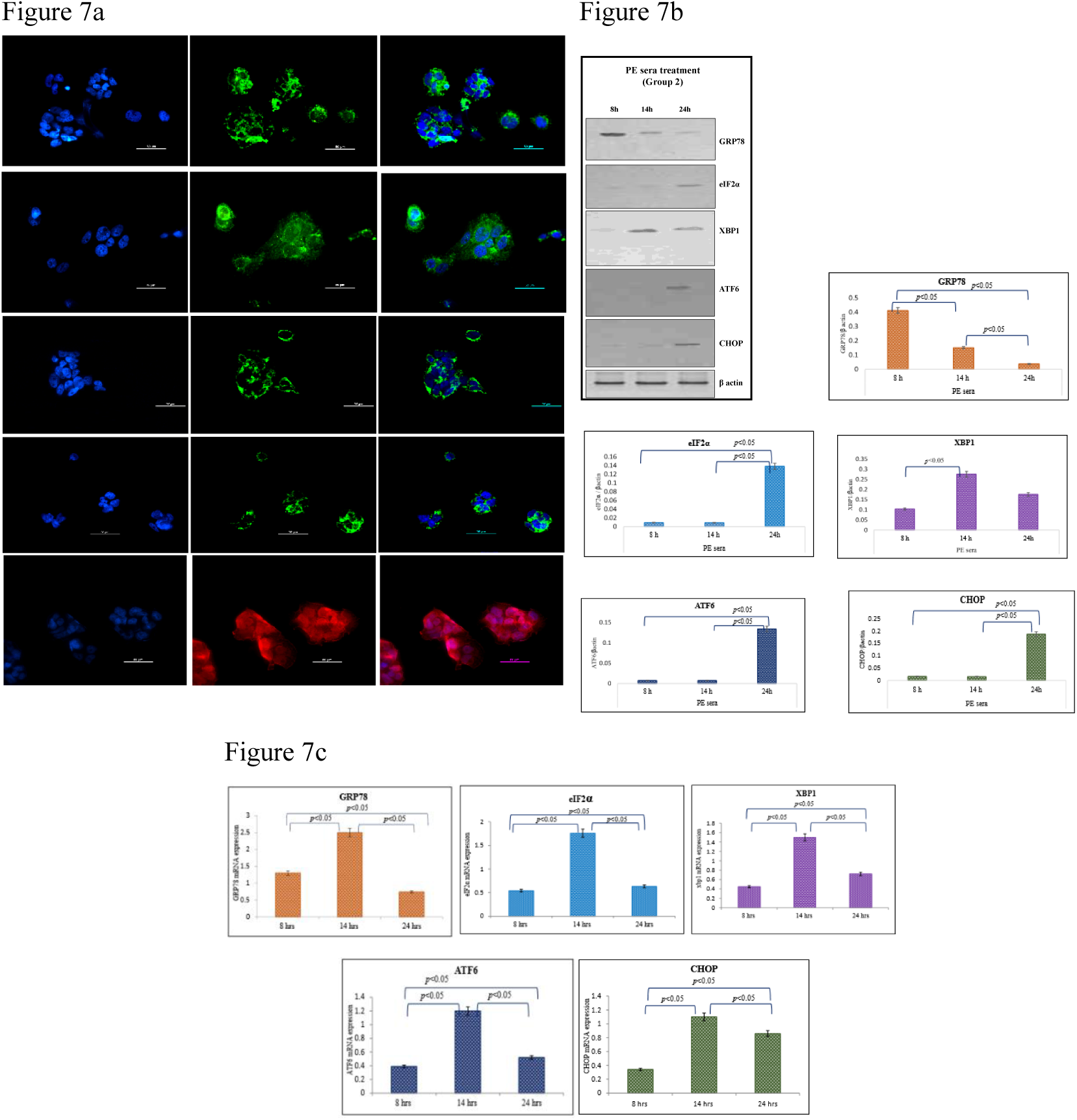
Representative immunofluorescence staining pattern of anti-GRP78 antibody positive BeWo cells following 8 h, anti-eIF2α, anti-XBP1, anti-ATF6 antibody positive BeWo cells following 14 h and anti-DDIT3 (CHOP) antibody positive BeWo cells following 24 h treatment of PE sera; Figure 7b: Representative images of immunoblot showing the expression of ER stress markers, GRP78, eIF2α, XBP1, ATF6 and CHOP in BeWo cells. β-Actin was used as protein loading control. The Bar diagrams represent the normalized values of the markers. Results are representative of 7 independent experiments. Data presented as mean ± SD. Statistical analysis was done using one way ANOVA with Bonferroni’s post hoc; Figure 7c: Bar diagrams represent the relative mRNA expression of GRP78, eIF2α, XBP1, ATF6 and CHOP and was found maximum at 14 h. GAPDH was used as positive control. Data presented as mean ± SD. One way ANOVA with Bonferroni’s post hoc test was applied (*p* values indicated on graph itself)

#### Expression/Intensity of ER stress markers in BeWo cells reduced when recombinant VEGF (re-VEGF) was added to sera from preeclamptic (PE) pregnant women [Figure 8a-c]

Immunofluorescence microscopy revealed that expressions of GRP 78, eIF2α, XBP1, ATF6 and CHOP were reduced significantly at 8, 14 and 24 hours [Figure 8a]. By immunoblot analysis, GRP78, eIF2α and CHOP expressions were low at 8 h and 14 h [Figure 8b]. mRNA levels were consistent with the protein expression [Figure 8c].

**Figure 8a:**
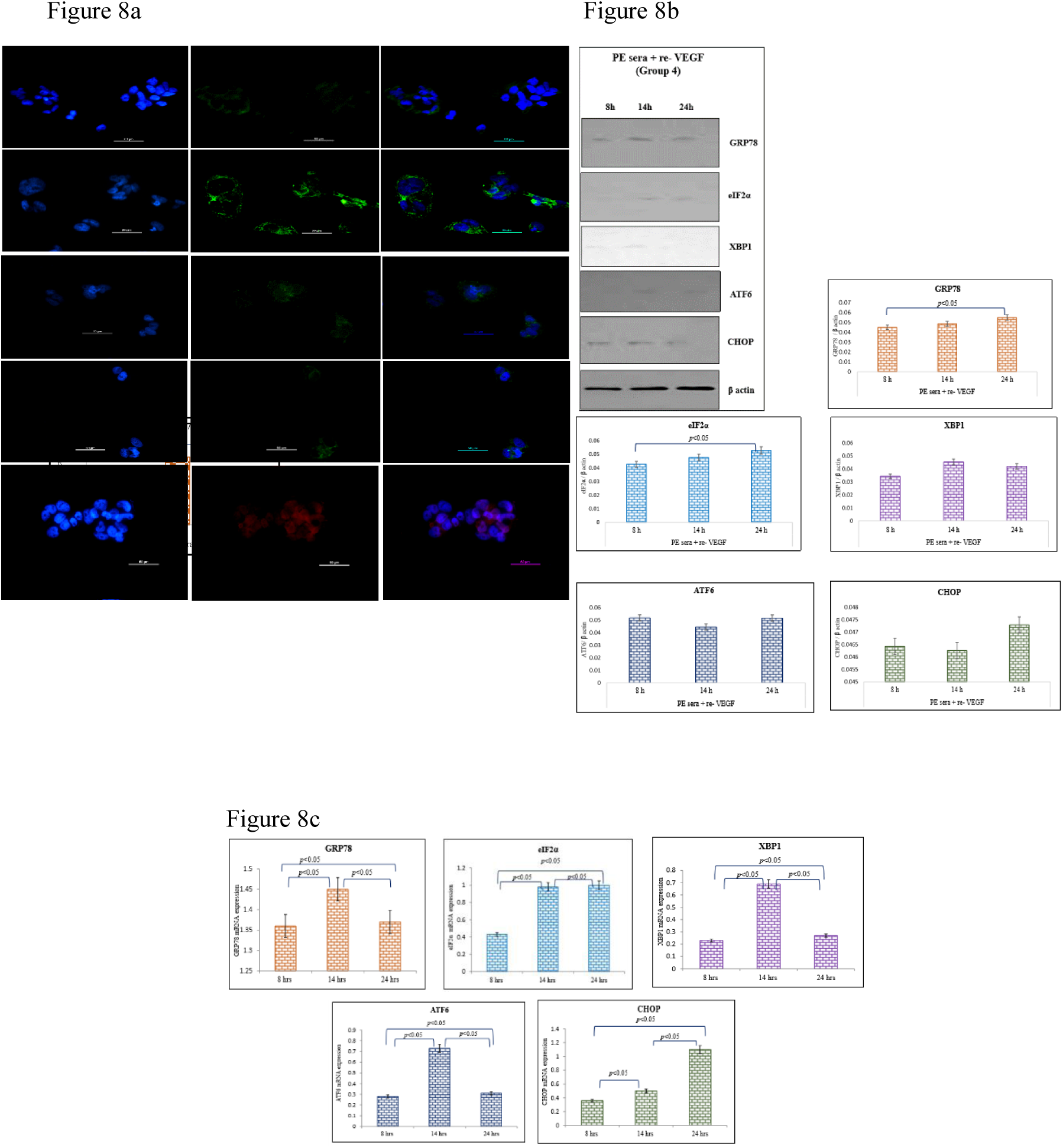
Representative immunofluorescence staining pattern of anti-GRP78 antibody positive BeWo cells following 8 h, anti-eIF2α, anti-XBP1, anti-ATF6 antibody positive BeWo cells following 14 h and anti-DDIT3 (CHOP) antibody positive BeWo cells following 24 h treatment of PE + re-VEGF sera; Figure 8b: Representative images of immunoblot showing the expression of ER stress markers, GRP78, eIF2α, XBP1, ATF6 and CHOP in BeWo cells. β-Actin was used as protein loading control. The Bar diagrams represent the normalized values of the markers. Results are representative of 7 independent experiments. Data presented as mean ± SD. Statistical analysis was done using one way ANOVA with Bonferroni’s post hoc; Figure 8c: Bar diagrams represent the relative mRNA expression of GRP78, eIF2α, XBP1, ATF6 and CHOP. GAPDH was used as positive control. Data presented as mean ± SD. One way ANOVA with Bonferroni’s post hoc test was applied (*p* values indicated on graph itself)

#### Raised expression of ER stress markers (GRP78, eIF2α, XBP1, ATF6 and CHOP) was noted in re-sFlt-1 (9 and 12 ng/ml) treated BeWo cells [Figure 9a-c]

Significant expression of XBP1, ATF6 and CHOP was observed at 14 hours with 12 ng/ml concentration of re-sFlt-1 as compared to 9 ng/ml.

ng/ml. Characteristic stress granules in case of eIF2α were also seen at 14 hours [Figure 9a]. Immunoblot revealed higher expressions of GRP78 and eIF2α at 14 h with 12 ng/ml concentration (c2) of recombinant sFlt-1 as compared to 9 ng/ml concentration (c1). Expression of XBP1 was

**Figure 9a:**
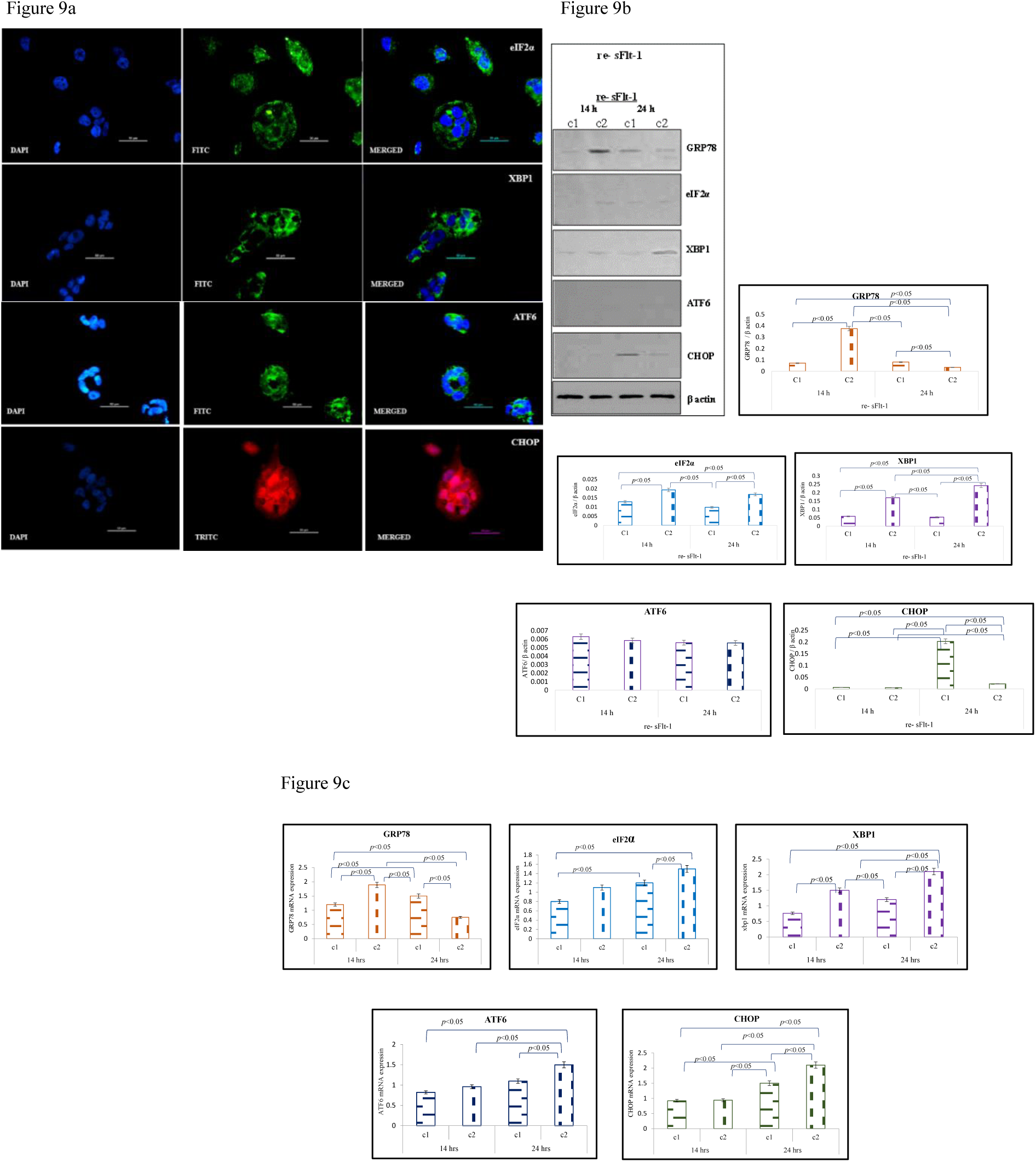
Representative immunofluorescence staining pattern of anti-eIF2α, anti-XBP1, anti-ATF6 and anti-DDIT3 (CHOP) antibody positive BeWo cells following 14 treatment of re-sFlt-1 (12ng/ml); Figure 9b: Representative images of immunoblot showing the expression of ER stress markers, GRP78, eIF2α, XBP1, ATF6 and CHOP in BeWo cells. β-Actin was used as protein loading control. The Bar diagrams represent the normalized values of the markers. Results are representative of 7 independent experiments. Data presented as mean ± SD. Statistical analysis was done using one way ANOVA with Bonferroni’s post hoc; Figure 9c: Bar diagrams represent the relative mRNA expression of GRP78, eIF2α, XBP1, ATF6 and CHOP. GAPDH was used as positive control. Data presented as mean ± SD. One way ANOVA with Bonferroni’s post hoc test was applied (*p* values indicated on graph itself)

found to be higher at 24 h with 12 ng/ml concentration (c2). ATF6 expression was found consistent at both time points (14 h and 24 h). CHOP expression was observed maximum at 24 h with 9 ng/ml (c1) concentration of recombinant sFlt-1 [Figure 9b]. GRP78 mRNA levels were found maximum at 14 h (c2) whereas mRNA levels of eIF2α, XBP1, ATF6 and CHOP were found maximum at 24 h (c2) [Figure 9c].

#### Earmarked expression of GRP78, eIF2α, XBP1, ATF6 and CHOP in Tunicamycin treated BeWo cells [Figure 10a-c]

The expression of all the markers were observed more so with 5 µg/ml dose of tunicamycin as compared to its lower dose [Figure 10a]. Immunoblot data revealed higher expressions of GRP 78 and eIF2α at 14h and 24 h as compared to 8 h. Expression was even more when its concentration was increased. XBP1, ATF6 and CHOP expressions was found to be higher with 5 µg/ml of tunicamycin as compared to 2.5 µg/ml of tunicamycin [Figure 10b]. GRP78, eIF2α, XBP1 and ATF6 mRNA levels were found to be higher at 14 h as compared to 8 h and 24 h. CHOP mRNA levels were found maximum at 24 h [Figure 10c].

**Figure 10a:**
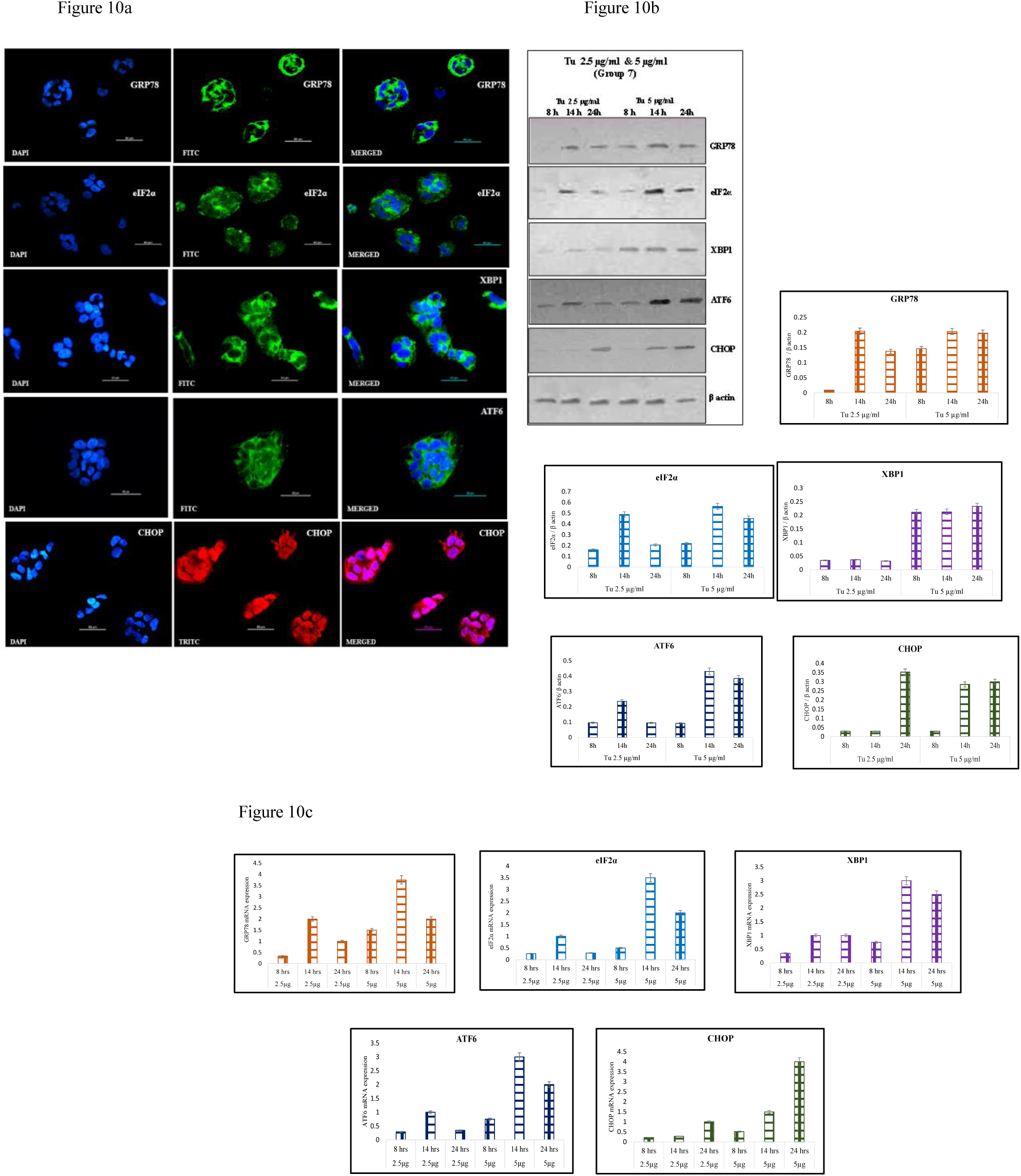
Representative immunofluorescence staining pattern of anti-GRP78 antibody positive BeWo cells following 8 h, anti-eIF2α, anti-XBP1, anti-ATF6 antibody positive BeWo cells following 14 h and anti-DDIT3 (CHOP) antibody positive BeWo cells following tunicamycin treatment (5µg/ml). Figure 10b: Representative images of immunoblot showing the expression of ER stress markers, GRP78, eIF2α, XBP1, ATF6 and CHOP in BeWo cells. β-Actin was used as protein loading control. The Bar diagrams represent the normalized values of the markers. Results are representative of 7 independent experiments. Data presented as mean ± SD. Statistical analysis was done using one way ANOVA with Bonferroni’s post hoc; Figure 10c: Bar diagrams represent the relative mRNA expression of GRP78, eIF2α, XBP1, ATF6 and CHOP. GAPDH was used as positive control. Data presented as mean ± SD. One way ANOVA with Bonferroni’s post hoc test was applied (*p* values indicated on graph itself)

#### Feeble expression of GRP78, eIF2α, XBP1, ATF6 and CHOP in untreated BeWo cells [Figure 11a-c]

Mild expression of GRP78 at 8 hours and eIF2α at 14 hours was observed. However, no expression of XBP1, ATF6 and CHOP was observed at different time points [Figure 11a]. Western blot analysis revealed higher expression of GRP78, eIF2α and XBP1 at 14 h as compared to 8 h and 24 h. ATF6 and CHOP expressions were found maximum at 24 h [Figure 11b]. GRP78, eIF2α and ATF6 mRNA levels were found maximum at 14 h however XBP1 and CHOP mRNA levels were found to be higher at 24 h [Figure 11c].

**Figure 11a:**
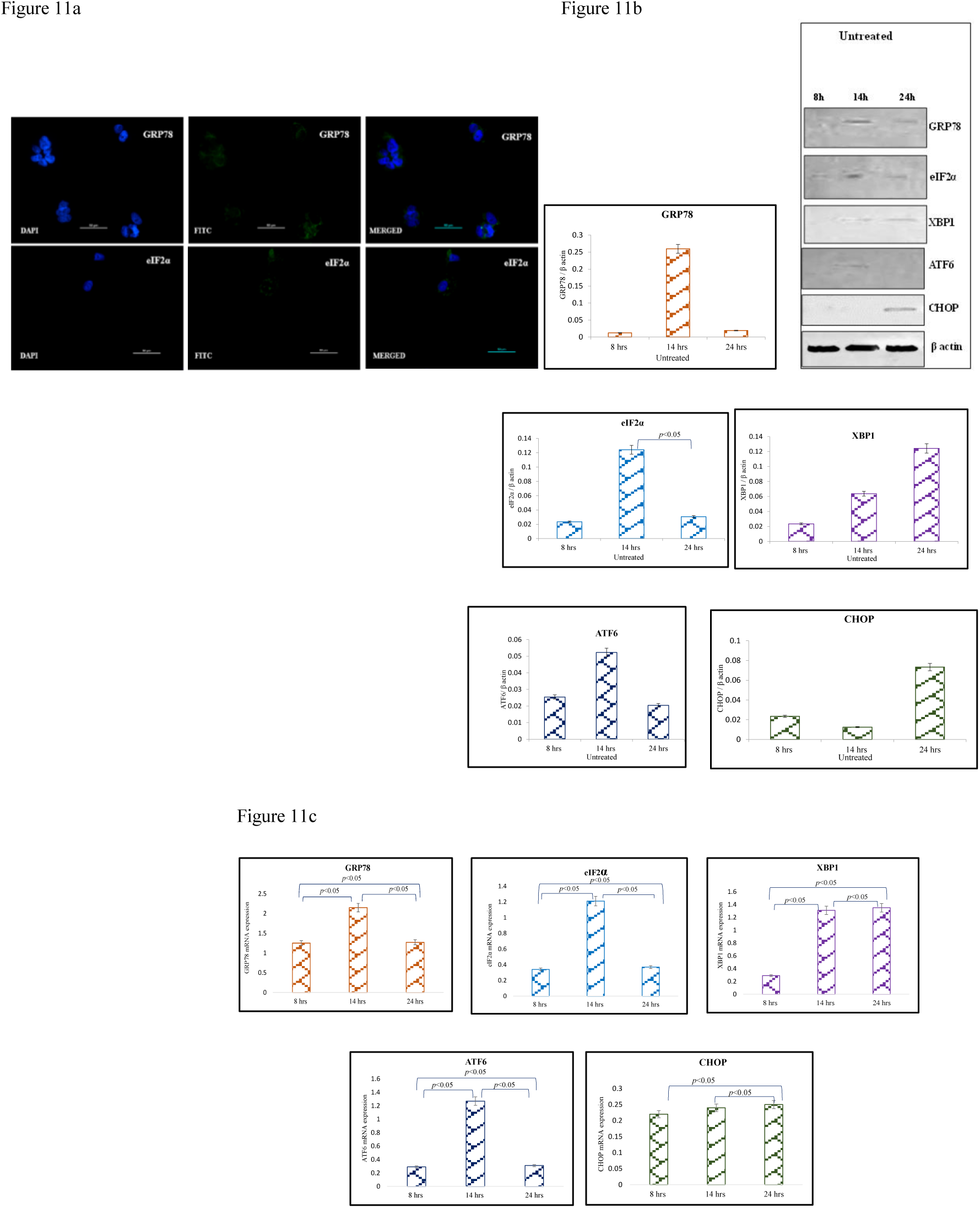
Representative immunofluorescence staining pattern of anti-GRP78 antibody positive BeWo cells following 8 h and anti-eIF2α antibody positive BeWo cells following no treatment. Figure 11b: Representative images of immunoblot showing the expression of ER stress markers, GRP78, eIF2α, XBP1, ATF6 and CHOP in BeWo cells. β-Actin was used as protein loading control. The Bar diagrams represent the normalized values of the markers. Results are representative of 7 independent experiments. Data presented as mean ± SD. Statistical analysis was done using one way ANOVA with Bonferroni’s post hoc; Figure 11c: Bar diagrams represent the relative mRNA expression of GRP78, eIF2α, XBP1, ATF6 and CHOP. GAPDH was used as positive control. Data presented as mean ± SD. One way ANOVA with Bonferroni’s post hoc test was applied (*p* values indicated on graph itself)

#### Comparison of normalized protein values and mRNA levels of various ER stress markers between NT sera and NT sera + re-sFlt-1 treated BeWo cells [Figure 12a-j]

At 8 h, expressions of all ER stress markers were found to be higher in NT+ re-sFlt-1 treated BeWo cells as compared to NT sera and the difference was found to be statistically significant [GRP78 (*p*<0.0001), eIF2α (*p*<0.0001), XBP1 (*p*<0.0001), ATF6 (*p*<0.0001) and CHOP (*p*<0.0001)] [Figure 12a-e]. At 14 h, GRP78, eIF2α, XBP1 and CHOP expressions were also higher in NT+ re-sFlt-1 treated BeWo cells as compared to NT sera and the difference was statistically significant [GRP78 (*p*<0.0001), eIF2α (*p*<0.0001), XBP1 (*p*<0.0001) and CHOP (*p*<0.0001)] [Figure 12a-e]. At 24 h, expressions of all ER stress markers were found to be higher in NT+ re-sFlt-1 treated BeWo cells as compared to NT sera treated BeWo cells and the difference was statistically significant. GRP78 (*p*<0.0001), eIF2α (*p*=0.0002), XBP1 (*p*=0.0129), ATF6 (*p*=0.0001) and CHOP (*p*<0.0001) [Figure 12a-e]. At 8, 14 and 24 h, GRP 78, eIF2α, XBP1, ATF6 and CHOP mRNA levels were found to be higher in NT+ re-sFlt-1 treated BeWo cells as compared to NT sera treated cells and the difference was found to be statistically significant [GRP78 (*p*<0.0001), eIF2α (*p*<0.0001), XBP1 (*p*<0.0001), ATF6 (*p*<0.0001) and CHOP (*p*<0.0001)] [Figure 12f-j].

**Figure 12a-e:**
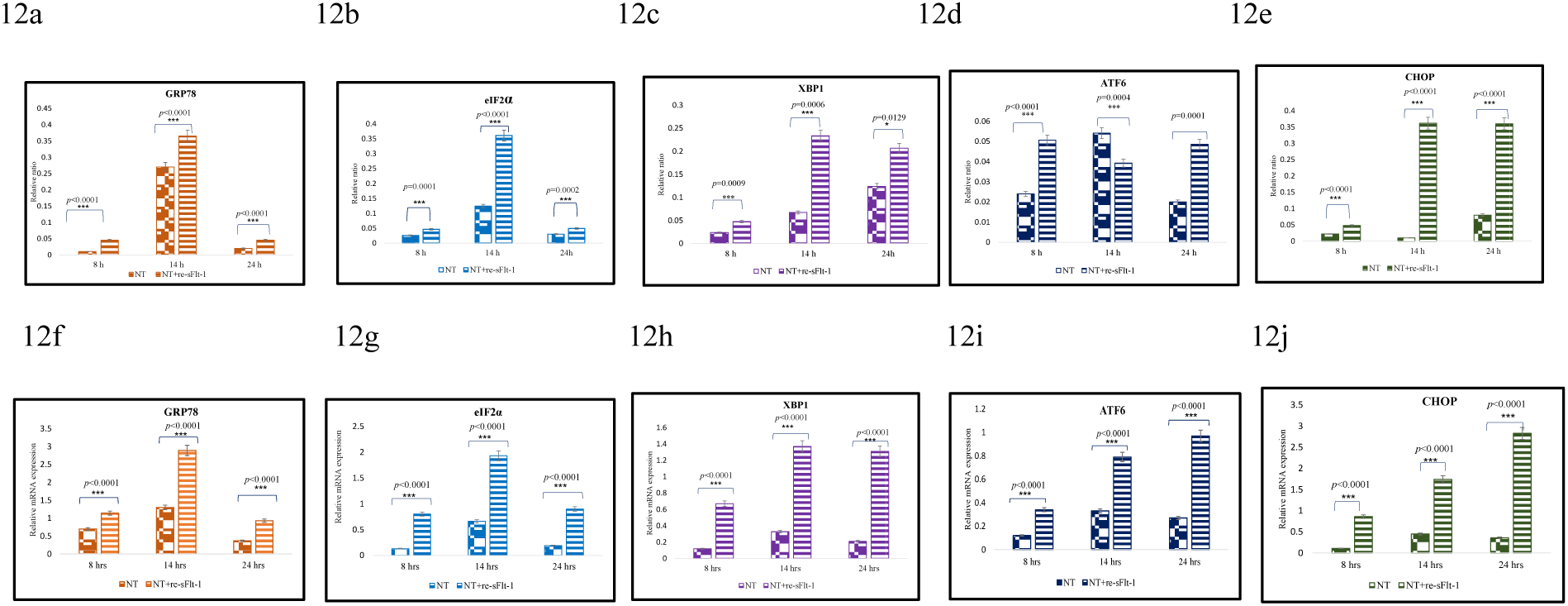
Comparison of normalized protein values of various ER stress markers between NT sera treated BeWo cells and cells which received both NT sera + re-sFlt-1; Figure 12f-j: Comparison of relative mRNA expression of ER stress markers between NT sera treated BeWo cells and cells which received both NT sera + re-sFlt-1. Statistical analysis was done using paired t test at all the indicated time points for all markers

#### Comparison of normalized protein values and mRNA levels of various ER stress markers between NT sera treated and PE sera treated BeWo cells [Figure 13a-j]

At 8 h, GRP78 and XBP1 expressions were found to be higher in PE sera treated BeWo cells as compared to NT sera treated cells and the difference was found to be statistically significant [GRP78 (*p*=0.0156) and XBP1 (*p*=0.0156)] [Figure 13a, 13c]. At 14 h, XBP1 and CHOP expressions were also higher in PE sera treated as compared to NT sera treated cells and the difference between two groups was statistically significant. [XBP1 (*p*<0.0001) and CHOP (*p*=0.0002)] [Figure 13c, 13e]. At 24 h, expressions of all ER stress markers were found to be higher in PE sera treated as compared to NT sera treated cells and the difference was statistically significant [GRP78 (*p*<0.0001), eIF2α (*p*<0.0001), XBP1 (*p*<0.0001), ATF6 (*p*<0.0001) and CHOP (*p*<0.0001)] [Figure 13a-e]. At 8, 14 and 24 h, GRP 78, eIF2α, XBP1, ATF6 and CHOP mRNA levels were found to be higher in PE sera treated as compared to NT sera treated BeWo cells and the difference was found to be statistically significant [GRP78 (*p*<0.0001), eIF2α (*p*<0.0001), XBP1 (*p*<0.0001), ATF6 (*p*<0.0001) and CHOP (*p*<0.0001)] [Figure 13f-j].

**Figure 13a-e:**
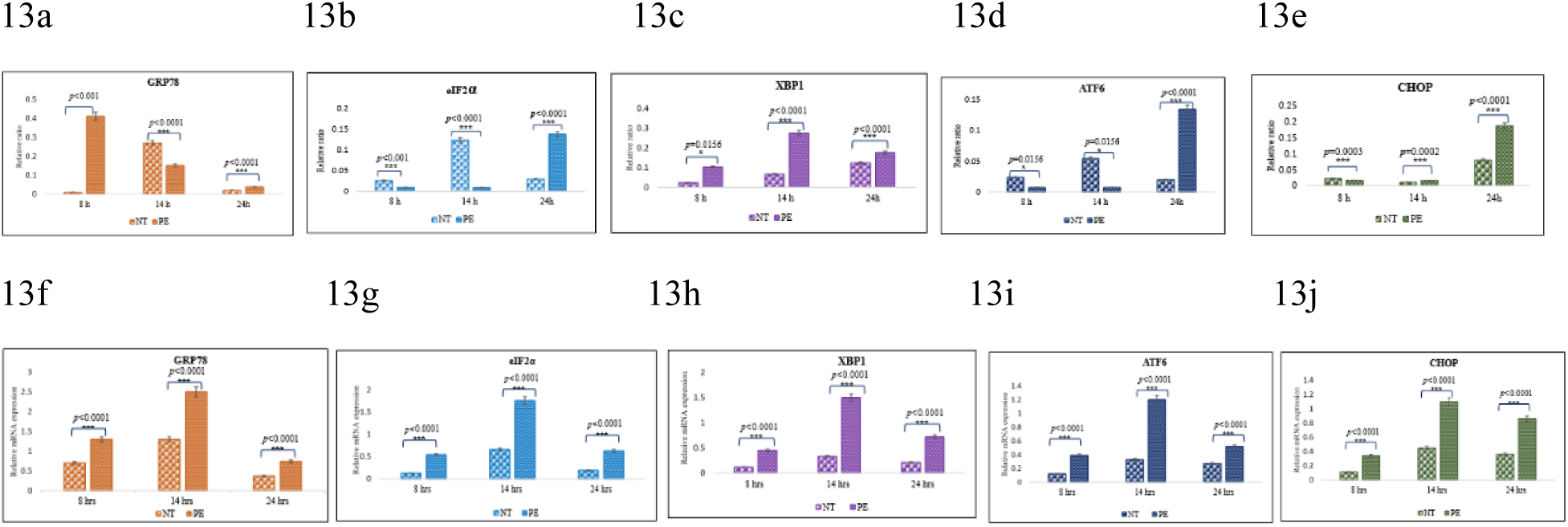
Comparison of normalized protein values of various ER stress markers between NT and PE sera treated BeWo cells; Figure 13f-j: Comparison of relative mRNA expression of ER stress markers between NT and PE sera treated BeWo cells. Statistical analysis was done using paired t test at all the indicated time points for all markers

#### Comparison of normalized protein values and mRNA levels of various ER stress markers between PE sera and PE sera along with recombinant VEGF (PE + re-VEGF) treated BeWo cells [Figure 14a-j]

At 8 h, GRP78 and XBP1 expressions were found to be lower in PE+ re-VEGF treated BeWo cells as compared to PE sera treated cells and the difference was found to be statistically significant GRP78 (*p*<0.0001) and XBP1 (*p*=0.0156) [Figure 14a, 14c]. At 14 h, GRP78 and XBP1 expressions were found to be lower in PE+ re-VEGF treated BeWo cells as compared to PE sera treated cells and the difference was statistically significant. [GRP78 (*p*<0.0001) and XBP1 (*p*<0.0001)] [Figure 14a, 14c]. At 24 h, expressions of eIF2α, XBP1, ATF6 and CHOP were found to be lower in PE+ re-VEGF treated BeWo cells as compared to PE sera treated cells and the difference was statistically significant. eIF2α (*p*<0.0001), XBP1 (*p*<0.0001), ATF6 (*p*<0.0001) and CHOP (*p*<0.0001) [Figure 14 b-e]. At 8 h, eIF2α, XBP1 and ATF6 mRNA levels were found to be lower in PE+re-VEGF treated BeWo cells as compared to PE sera treated cells and the difference was found to be statistically significant [eIF2α (*p*<0.0001), XBP1 (*p*<0.0001), ATF6 (*p*<0.0001)] [Figure 14g-i]. At 14 h, GRP78, eIF2α, XBP1, ATF6 and CHOP mRNA levels were found to be lower in PE+ re-VEGF treated BeWo cells as compared to PE sera treated cells and the difference was found to be statistically significant [GRP78 (*p*<0.0001), eIF2α (*p*<0.0001), XBP1 (*p*<0.0001), ATF6 (*p*<0.0001) and CHOP (*p*<0.0001)] [Figure 14f-j]. At 24 h, XBP1 and ATF6 mRNA levels were found to be lower in PE+ re-VEGF treated BeWo cells as compared to PE sera treated cells and the difference was found to be statistically significant [XBP1 (*p*<0.0001), ATF6 (*p*<0.0001)] [Figure 14h-i].

**Figure 14a-e:**
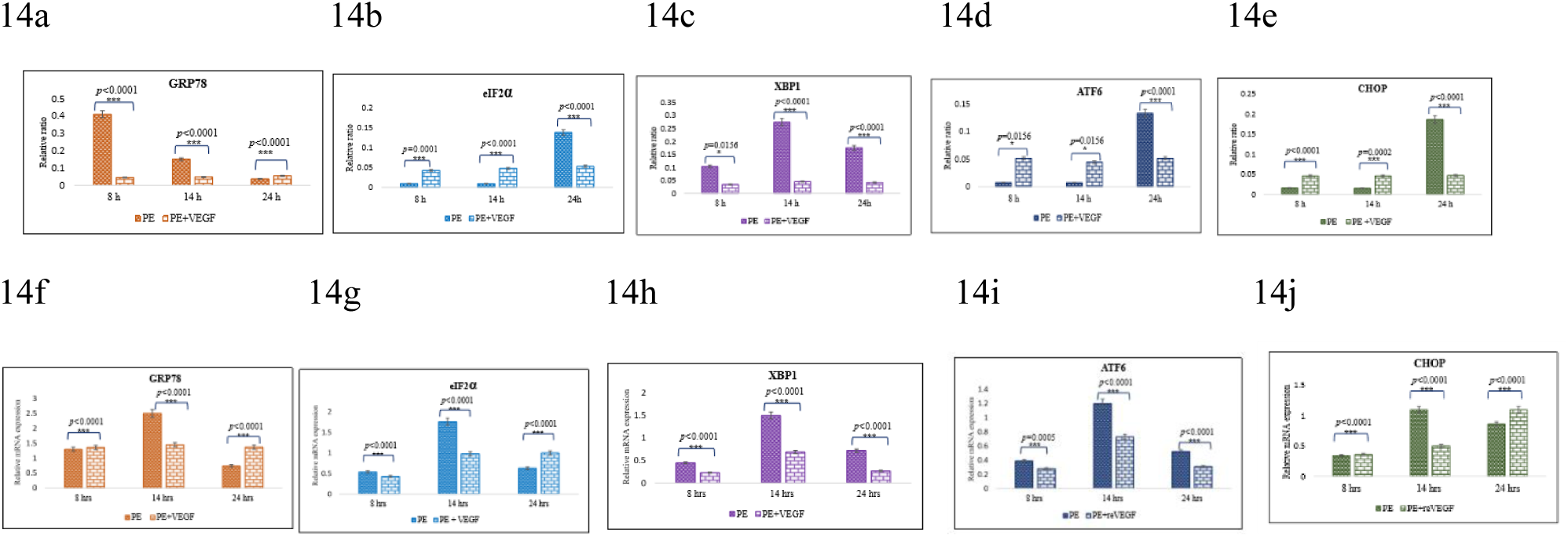
Comparison of normalized protein values of various ER stress markers between PE sera and PE sera and recombinant VEGF (PE + re-VEGF) treated BeWo cells; Figure 14f-j: Comparison of relative mRNA expression of ER stress markers between PE sera and PE sera and recombinant VEGF (PE + re-VEGF) treated BeWo cells. Statistical analysis was done using paired t test at all the indicated time points for all markers

## Discussion

In the present study we reported reduced serum levels of VEGF and increased level of sFlt-1 in women with preeclampsia suggesting an anti-angiogenic state may be involved in the pathogenesis of PE**^21^**. Angiogenesis, the formation of new blood vessels from pre-existing ones, is essential for a successful pregnancy. The role of VEGF as a key regulator of normal and abnormal angiogenesis has already been established over the years**^22^**. VEGF stabilizes endothelial cells in mature blood vessels, crucial for maintaining endothelial cell homeostasis. It promotes neovascularization, contributes to maintenance of vascular tone by influencing NO production, reduces blood pressure and is important for maintenance of normal glomerular filtration**^23,24,25^**. Therefore, low levels of VEGF could cause disturbances in the placental development and affect maternal vessels as seen in PE. Under normal physiological conditions, VEGF exerts its biological effects through two high affinity tyrosine kinase receptors VEGFR1 or fms like protein kinase-1/Flt-1 and VEGFR2 or kinase insert domain receptor/KDR**^26,27^**. The soluble isoform of VEGFR1 (Flt-1) known as s-VEGFR1(sFlt-1), binds with free VEGF in circulation with higher affinity thereby preventing its interaction with its endogenous receptors (VEGFR1 and VEGFR2) and thus reducing its biological effects like placental development and maintenance of maternal vasculature. Thus, the reduced free VEGF due to increased sFlt-1 (imbalance of pro and anti-angiogenic factors) could alter the balance of superoxide anion and nitric oxide, resulting in increased oxidative stress contributing to vascular endothelial dysfunction**^28^.** Although, it is known that both stress processes (oxidative stress and ER stress) are intimately related, the mechanisms linking them are not fully understood, however these two may be linked by three principal ways**^29^**. Placental endoplasmic reticulum stress has recently been recognized in the pathophysiology of preeclampsia. The vulnerability of placental cells i.e., syncytiotrophoblast to ER stress may be due to their involvement in extensive secretory activity. Also molecular and morphological evidence confirms high levels of ER stress in placentas from cases of early onset of preeclampsia as compared to late onset**^30^**.

After the onset of ER stress (on failure of proteostasis), initial event is the dissociation of GRP78 from trans-membrane sensors, PERK (PKR-like ER kinase), IRE1 (inositol-requiring enzyme 1) and ATF6 (activated transcription factor 6). These sensors are integral ER membrane proteins that signal luminal stress to the cytosol and the nucleus. When ER stress is not mitigated and homeostasis is not restored, the UPR(unfolded protein response) triggers apoptosis. The mammalian UPR, in chronological time frame has evolved into a network of signaling events that responds to various inputs over a wide range of basal metabolic states. A number of conditions can activate the UPR, including changes in glycosylation, redox status, glucose availability, calcium homeostasis and secretory protein load**^31^**.

Thus in the present study we hypothesized that levels of GRP78 are raised in sera of patients of PE. To validate this hypothesis, we estimated the GRP78 levels in serum by sandwich ELISA. The maternal serum levels of GRP78 were found to be higher in PE patients as compared to normotensive, non-proteinuric pregnant women (table 4). With this finding, we deciphered that the raised levels of GRP78 in patients’ sera might be the reason for its earlier dissociation from trans-membrane sensors and its subsequent release in the maternal serum thereby inciting ER stress following signaling through individual ER stress arms. However these results did not establish a cause and effect relationship i.e. whether the sFlt-1 is the causative factor for increased GRP78 found in PE patients. In order to answer this question, our group for the first time performed *in-vitro* experiments using BeWo cells which received various treatments having varying concentration of sFlt-1 (normotensive and preeclamptic sera) and the expression of ER stress markers were analyzed at transcript and protein level at different time points. The three different time points of study (8h,14h and 24 h) were chosen to observe the activation and expression of markers after activation of ER stress pathway (confirmed after the signal from master regulator of UPR) up to the signal of apoptotic marker (CHOP).

The prime regulator of UPR *i.e.* GRP78, in the resting condition, binds to N-terminus of trans-membrane sensors, but its dissociation and subsequent competitive binding to accumulating misfolded proteins leads to dimerization, autophosphorylation and activation of PERK and IRE1. The expression of GRP78 transcript and protein was detected as early as 8h however maximum expression varied from 14 to 24 h in various groups. The levels of both GRP78 transcripts and protein were further upregulated whenever BeWo cells were treated with sera having high sFlt-1 concentration as in group 2(NT+ re-sFlt-1), 3(PE sera) and 5(recombinant s-Flt-1) [figure 6a-c, 7a-c, 9a-c]. In contrast, significantly lower expression of GRP78 was noted when BeWo cells got exposed to preeclamptic sera along with recombinant VEGF (group 4:PE+re-VEGF) as compared to PE sera alone treated BeWo cells (group 3:PE) at both protein as well as transcript levels. Our results are in concurrence with the evidences in literature demonstrating sFlt-1 as a potent inhibitor of VEGF indicating that the addition of re-VEGF neutralized the sFlt-1 present in PE sera**^32^**. The upregulated expression of GRP78 in group 2 and group 3 thus suggested the presence of a common factor which might increase the expression of GRP78 (and in turn ER stress) at both protein and mRNA levels. We inferred, this factor might be recombinant sFlt-1 in group 2 and increased concentration of sFlt-1 in group 3. However decrease in the expression of GRP78 in group 4 indicates neutralization of increased concentration of sFlt-1 by recombinant VEGF (re-VEGF).

Emerging evidence indicates amplitude and kinetics of UPR signaling are tightly regulated at different levels, which has direct impact on cell fate decisions. How protein folding stress at the ER is sensed has been a central topic in the research field for more than a decade. Different models have been proposed to explain how ER stress is sensed, and these are constantly modified over time owing to new findings and to discrepancies and similarities between the yeast and mammalian UPR**^33^**. Activation of PERK results in the phosphorylation of eukaryotic initiation factor 2 subunit α (eIF2 α), blocking protein translation and reducing the protein burden within the ER. Adaptive response goes as long as the level of stress is below threshold, however in case of prolonged stress, eIF2α-ATF4-CHOP pathway gets activated, it drives the cells toward apoptosis. In the present study, expression of eIF2α was observed maximum at 14 h both in group1 and 2 however more in group 2(addition of re-sFlt-1 to NT sera) at both protein and mRNA levels. Delayed attenuation of PERK arm and concomitant signal at 24 h was reported in BeWo cells treated with high sFlt-1 concentration (group 3). Our finding is consistent with the study carried out by Walter *et. al.* (2007) where they observed eIF2α signal till 30 h**^34^**. In contrast, as we expected, addition of re-VEGF to PE sera (group 4) reduced (because of neutralization of sFlt-1 by re-VEGF) the eIF2α expression at all the time points. The mRNA levels were found consistent with the protein expression [Figure 8a-c].

IRE1α was identified as a positive regulator for cell survival. It was believed that IRE1α signaling was terminated during irremediable ER stress to enable apoptosis**^35,36,37,38^**. A dissociation of IRE1α from GRP78 (BiP) due to an elevated level of unfolded proteins in the ER leads to activation of IRE1α. As an ER transmembrane protein, it monitors ER homeostasis through an ER luminal stress sensing domain and triggers UPR through a cytoplasmic kinase domain and an RNase domain**^39^**. Upon ER stress, IRE1 RNase is activated through conformational change, autophosphorylation, and higher order oligomerization**^40,41,42^**. While IRE1 promotes cell survival, it can also initiate apoptosis via decay of anti-apoptotic microRNAs**^43,44^**. It initiates diverse downstream signaling of the UPR either through unconventional splicing of the transcription factor XBP-1 or through post transcriptional modifications via Regulated IRE1-Dependent Decay (RIDD) of multiple substrates**^45,46,47,48^**. The spliced XBP-1 enters into the nucleus to transcriptionally reprogram UPR target genes. We observed that protein and mRNA expression of XBP1 was observed maximum at 14 h in cells treated with high sFlt-1 concentration (BeWo cells exposed with PE sera) which is consistent with the previous study by walter *et. al.* in 2007**^49^**. The above pattern observed in IRE1 arm was similar to that of PERK arm in the studied groups and the noted difference in the expression of markers were significant whether it was between group 1and 3 or group 3 and 4 [Figure 13c, 14c, 14 h]. Mammals possess two homologous ATF6 proteins, ATF6α and ATF6β**^50^**. ATF6α participates in the induction of various UPR target genes. ATF6β seems to have a minimal role in the UPR**^51,52^**. In response to ER stress, ATF6 translocates from the ER to the Golgi where it is first cleaved by site-1 protease (S1P) in its luminal domain, separating ATF6 into two halves. The NH2-terminal half remains anchored to the membrane and is subsequently cleaved by site-2 protease (S2P). The cytosolic domain of ATF6 is then liberated from the membrane and translocates to the nucleus and induces expression of genes with ER stress response element (ERSE) in their promoter such as the ER-chaperone protein BiP and the transcription factors C/EBP homologous protein (CHOP) and X-box binding protein-1 (XBP1)**^53^**. In the present study, ATF6 expression was also found to be increased in BeWo cells of group 2 and 3 as compared to group 1 and 4 at both protein and mRNA levels [figure 12d, 12 i, 14d, 14i]. The maximum expression was noted between 14 to 24 h.

If the various UPR-induced mechanisms fail to alleviate ER stress, both the intrinsic and extrinsic pathways for apoptosis can become activated. Players involved in the cell death response include (i) PERK/eIF2α-dependent induction of the pro-apoptotic transcriptional factor CHOP; (ii) IRE1-mediated activation of TRAF2 (tumor necrosis factor receptor associated factor 2), which stimulates the ASK1 (apoptosis signal-regulating kinase 1)/JNK (c-Junamino terminal kinase) kinase cascade, and (iii) Bax/Bcl2-regulated Ca2+ release from the ER. CHOP/GADD153 (growth arrest/DNA damage) plays a convergent role in the UPR and it has been identified as one of the most important mediators of ER stress-induced apoptosis protein**^54^**. CHOP (CCAAT-enhancer-binding *protein* homologous *protein*) has been shown to be involved in ER stress-induced apoptosis both *in vitro* and *in vivo*. CHOP-deficient MEFs were partially protected against ER stress-induced apoptosis, and kidneys of CHOP−/−mice showed less apoptosis after treatment with the ER stressor tunicamycin, an inhibitor of glycosylation**^55^**. The levels of both CHOP transcripts and protein were upregulated whenever BeWo cells were treated with sera having high sFlt-1 concentration as in group 2, 3 [figure 6a-c, 7a-c] and downregulated in group 1 and 4 [figure 5a-c, 8a-c] (lower concentrations of s-Flt-1). The maximum signal encountered at 24 h indicating induction of apoptotic machinery with progression of time. In the present study, BeWo cells, when treated with two independent concentrations of recombinant sFlt-1 only (group5), no significant expression of ER stress markers were seen either at protein or mRNA levels at 8 h however at 14 h, upregulation in all the ER stress markers except ATF6 and CHOP were noticed with 12ng/ml. Our current study highlights the intensity of expression of all the markers in the time frame of 8 to 24 h. These observations suggested that exogenous addition of recombinant sFlt-1 only was not sufficient enough to induce the ER stress at 8 h and it may require the assistance of some additional factors as may be present in PE serum. Miyaki *et.al.*(2016) have shown decrease in placental weight in mice whose cells overexpressed human s-Flt-1. Later the same group examined the direct effect of recombinant sFlt-1 on two ovarian cancer cell lines, one colorectal cancer cell line (*in vitro study*) and nude mice with ovarian tumor model (*in vivo study*) and elaborated the mechanism underlying the anti-angiogenic or anti-tumor effects of sFlt-1**^56^**. They observed that reduction in tumor volume in mice model was correlated positively to the dose of re-sFlt-1. However, in the present study, we focused on the effects of sFlt-1 on ER stress and found that markers of ER stress were up-regulated though at different time points when BeWo cells were treated with high sFlt-1 concentration and their expression reduced on treatment with low sFlt-1 concentration. To the best of our knowledge, there is no literature reporting about the role of sFlt-1 in inducing ER stress in trophoblast cells.

### Summary and conclusion

In the present study we demonstrated endoplasmic reticulum stress in trophoblast cells (BeWo cells) which reasserted the setback of unfolded protein response aiming to maintain proteostasis of the cellular environment. The increased signaling response of ER stress makers in the BeWo cells exposed to increased sFlt-1 concentration (NT + re-sFlt-1 and PE sera treated BeWo cells) and also with the exogenous addition of recombinant sFlt-1 independently, gesticulated that sFlt-1 may be one of the various factor(s) responsible for endoplasmic reticulum stress seen in placental trophoblast cells of preeclamptic women. Conversely, the addition of recombinant VEGF to preeclamptic sera neutralized the increased concentration of sFlt-1 and demonstrated down-regulation in expression of ER stress markers at both protein and transcript levels. The raised serum GRP78 level observed in women with preeclampsia as compared to normotensive non-proteinuric pregnant women might be due to raised sFlt-1. Increased placental stress beyond threshold may be fatal to both mother and conceptus during pregnancy. Attempts to reduce placental stress therefore must be a research priority prospectively. Our study can serve as an experimental and therapeutic template to investigate newer therapeutic regime in the management of preeclampsia. In future, potential therapeutic interventions for preeclampsia must therefore be designed to address trophoblastic stress in its entirety, rather than particular stress response pathways. Moreover, to further validate our findings, different studies can be designed by using sFlt-1 transfection in trophoblast cells and also using primary cell cultures or explants from preeclamptic and normotensive and non-proteinuric placentae. Different drugs can be designed to modulate the activities of various ER sensors in order to alleviate ER stress. Thus, more studies are required to identify various pharmacological agents to counteract the effects of sFlt-1 as therapeutic options for preeclampsia. Possible approaches to antagonize sFlt-1 and its physiological effects may include saturating the system with its natural ligands VEGF and PLGF or administering the anti sFlt-1 antibodies or use of small interfering RNA to reduce sFlt-1. If such expedients prove effective in alleviating the ER stress and manifestation of the symptoms of preeclampsia; the delivery could be safely postponed for even a few weeks which would have significant impact on neonatal morbidity and mortality.

## Authors Contributions

SM and RD conceived and designed the study. SM did all the experiments. PA assisted in experiments. NB provided the clinical inputs and critical suggestions. SDD assisted in statistical analysis. SM and RD wrote first draft. MKD gave a hand in re-editing the content. KL, AS, RK, NR, SKG, and SS assisted in compilation of the final draft.

