## SUPPLEMENTARY FILE (TABLE) for "sFlt-1 commutes unfolded protein response into endoplasmic reticulum stress in trophoblast cells in preeclamptic pregnancies"

| Gene | Forward primer | Reverse primer |
|---|---|---|
| GRP78 | 5´-TGTTCAACCAATTATCAGCAAACTC - 3´ | 5´ - TTCTGCTGTATCCTCTTCACCAGT - 3´ |
| eIF2α | 5´ - AAGCATGCAGTCTCAGACCC - 3´ | 5´- GTGGGGTCAAGCGCCTATTA - 3´ |
| XBP1 | 5´ - TGGCCGGGTCTGCTGAGTCCG- 3´ | 5´ - ATCCATGGGGAGATGTTCTGG- 3´ |
| ATF6 | 5´ - CCACTAGTAGTATCAGCAGGAACTC- 3´ | 5´ - CCTTCTGCGGATGGCTTCAA- 3´ |
| CHOP | 5´ - AGAACCAGGAAACGGAAACAGA - 3´ | 5´ - TCTCCTTCATGCGCTGCTTT - 3´ |
| GAPDH | 5´ - AGCCGAGCCACATC - 3´ | 5´ - TGAGGCTGTTGTCATACTTCTC - 3´ |
| β-Actin | 5´ - GAGCACAGAGCCTCGCCTTT - 3´ | 5´ - TCATCATCCATGGTGAGCTGG - 3´ |

Table 1: Primers: Designed by NCBI

| Study Groups | | | |
|---|---|---|---|
| **Clinical characteristics** | Preeclampsia (n=30) | Normotensive, Non proteinuric (Control) (n=30) | Statistical significance (p value)* |
| Systolic blood pressure (mmHg) | 158.9 ±11.88 | 117.8±7.34 | p<0.0001 |
| Diastolic blood pressure (mmHg) | 101.43±8.39 | 74.2±6.39 | p<0.0001 |
| Body Mass Index | 27.83±5.61 | 23.67±3.59 | p<0.0001 |
| Protein (gm/day) | 4.9±1.6 | 0.7±0.2 | p<0.0001 |

**Table 2:** Clinical Characteristics of Preeclamptic and normotensive, non proteinuric pregnant women (controls) n= number of subjects, Data presented as mean±SD, Paired t test, *statistical significance, *p*<0.05

| Serum levels | Preeclampsia n=30 | Normotensive non proteinuric (Control) n=30 | Statistical significance (p Value)* |
|---|---|---|---|
| **sFlt-1** | 11295.25<br>2936.2 - 37818<br>Median (Range) | 2936.2<br>1180.43 - 6706.6<br>Median (Range) | 0.0001 |
| **VEGF** | 170.53 ± 36.55<br>Mean±SD | 254.61 ± 47.39<br>Mean±SD | 0.0001 |
| **GRP78** | 1103 ± 104.27<br>Mean±SD | 1018.61 ± 125.51<br>Mean±SD | 0.012 |

**Table 3:** Maternal serum levels of sFlt-1,VEGF and GRP78 in preeclamptic and normotensive, non proteinuric pregnant women (controls). n= number of subjects, Data presented as median (range), Paired t test, *statistical significance, *p*<0.05
